# Whole Genome Sequencing-based Characterization of Human Genome Variation and Mutation Burden in Botswana

**DOI:** 10.1101/2020.12.15.422821

**Authors:** Prisca K. Thami, Wonderful T. Choga, Delesa D. Mulisa, Collet Dandara, Andrey K. Shevchenko, Melvin M. Leteane, Vlad Novitsky, Stephen J. O’Brien, Myron Essex, Simani Gaseitsiwe, Emile R. Chimusa

## Abstract

The study of human genome variations can contribute towards understanding population diversity and the genetic aetiology of health-related traits. We sought to characterise human genomic variations of Botswana in order to assess diversity and elucidate mutation burden in the population using whole genome sequencing. Whole genome sequences of 390 unrelated individuals from Botswana were available for computational analysis. The sequences were mapped to the human reference genome GRCh38. Population joint variant calling was performed using Genome Analysis Tool Kit (GATK) and BCFTools. Variant characterisation was achieved by annotating the variants with a suite of databases in ANNOVAR and snpEFF. The genomic architecture of Botswana was delineated through principal component analysis, structure analysis and F_ST_. We identified a total of 27.7 million unique variants. Variant prioritisation revealed 24 damaging variants with the most damaging variants being *ACTRT2* rs3795263, *HOXD12* rs200302685, *ABCB5* rs111647033, *ATP8B4* rs77004004 and *ABCC12* rs113496237. We observed admixture of the Khoe-San, Niger-Congo and European ancestries in the population of Botswana, however population substructure was not observed. This exploration of whole genome sequences presents a comprehensive characterisation of human genomic variations in the population of Botswana and their potential in contributing to a deeper understanding of population diversity and health in Africa and the African diaspora.

## INTRODUCTION

Variants (or variations) are differences between the human reference genome (for instance GRCh38) and a genome of interest (1). Genomic variations (variants) include single nucleotide variants (SNVs), short insertions and deletions (indels) of less than 50 bases and structural variations (1–5). At the core of variant characterization is annotation of the discovered variants. Variant annotation involves interpreting the variants by determining their types, minor allele frequencies, effect prediction and predicting the genomic location of the discovered variants (6–9). Prioritizing variants in medical genetics mainly entails distinguishing background benign variants from pathogenic variants that can lead to disease phenotypes (10,11). According to comparative genetics, if a variant occurs in a gene that is conserved among species, this variant is likely to be pathogenic. In this regard a number of conservation methods can be used to identify deleterious mutations (12).

Characterizing genetic variations fosters the understanding of pathophysiology, and reconstruction of population histories through the inference of genetic relatedness (or divergence) of individuals (13,14). In genetic epidemiology, population structure is assessed and corrected to minimize spurious genetic associations and unmask signals of association (14–17). Patterns of genetic variation can be detected and quantified using methods such as Principal Components Analysis (PCA) (17,18), Wright’s fixation index (F_ST_) (19,20), patterns of homozygosity (21–23) and admixture proportion inference (24).

Major events such as the “Bantu expansion” and Eurasian migration into Southern Africa have shaped the genetic landscape of the region. These events have led to varying degrees of admixture of the migrant groups and indigenous population (14,25–31). Given the complex genetic architecture and high disease burden in Southern Africa, it is important to characterize the genetic variation within the region in order to understand the biology of disease (14,27,32).

We sought assess population genetic diversity and elucidate mutation burden in Botswana. Botswana is a landlocked country at the centre of Southern Africa (33). The population of Botswana is made up of mainly Bantu-speakers, an ethnolinguistic group of the Niger-Congo phylum (34–36). According to the latest population census the population is mainly comprised of about 92.7% Bantu-speakers, 1.7% Khoe-San, 3.6% Europeans, and relatively less Indians, and Asians (37). Since admixture occurs when previously isolated populations interbreed, it is possible to observe admixture of the aforementioned populations in Botswana. Using whole genome sequencing, we present a comprehensive characterization of the genomic variation and elucidate mutation burden within a Southern African population of Botswana.

## RESULTS

### Characterization of variants and variants effect

We identified 27.7 million variants from 390 individuals of Botswana. Of these variations, we found 25.1 million SNVs and 2.6 million indels (**Table S1**); 13.4% of these variations were novel, i.e. not found in dbSNP151, 1KGP, AGVP and gnomAD (38) (**Figure 1a**). The average transition-transversion (TI/TV) ratio was 2.1. The novel variants were classified into genomic region and functional classes. Of the 2,789,599 novel variants, intergenic variants were observed at the highest frequency (1,461,193), followed by intronic (1,066,166) and ncRNA (178,178) variants (**Figure 1b** and **Table S2**). A majority of the novel variants were singletons, rare (MAF <= 0.01) and low frequency variants (MAF >0.01-0.05) (**Table S3, Figure 1a** and **c**). Nonsynonymous SNVs, stop gain and stop loss variants formed 65.6% of the exonic variants (**Figure 1d** and **Table S2**).

**Figure 1.**
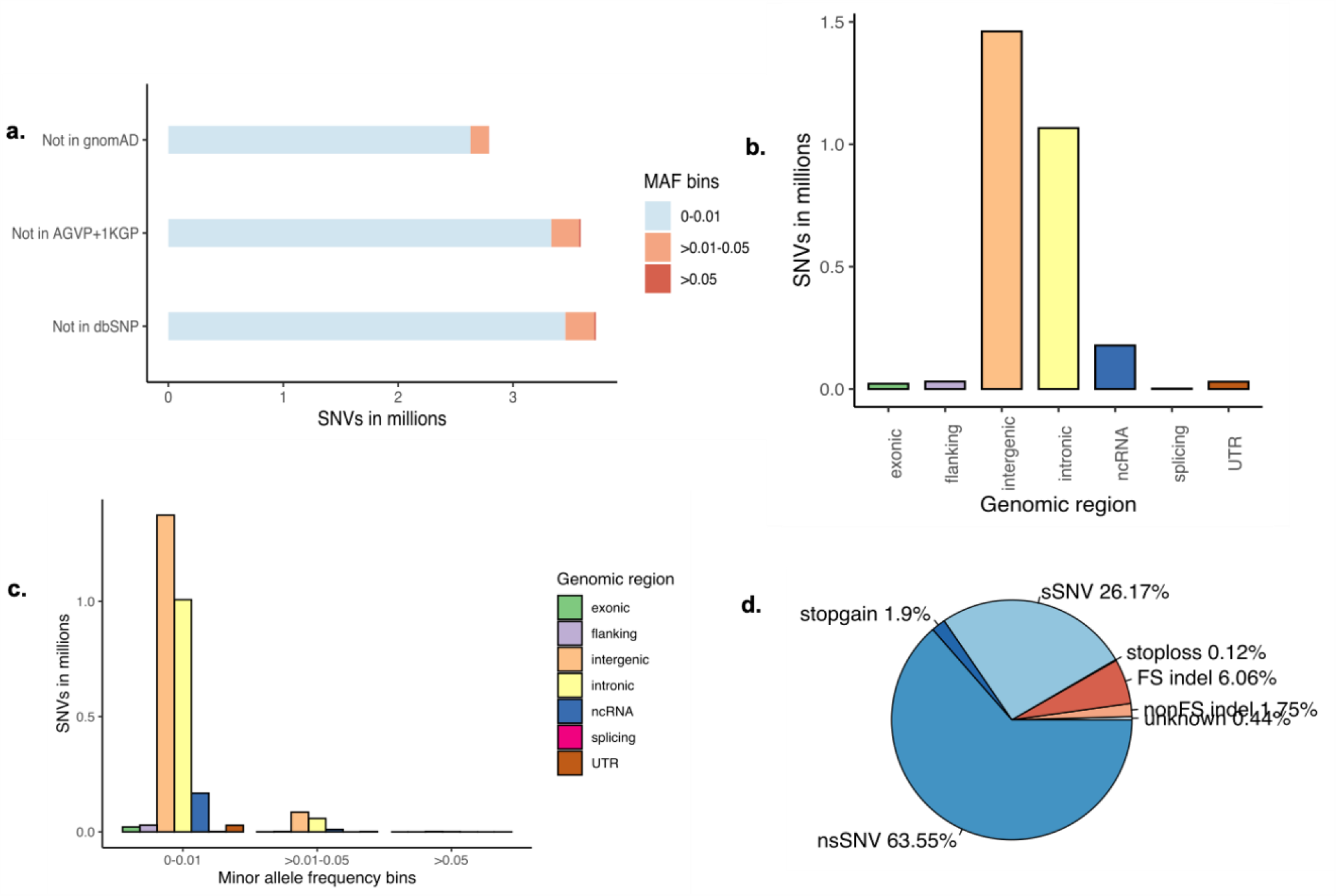
The distribution of novel variants in the Botswana population genomes. **a**. Novel variants, absent from dbSNP151, the African Genome Variation Project (AGVP), the 1000 Genomes Project (1KGP) and gnomAD. **b**. Genome-wide distribution of novel variant effects by functional elements. **c**. Distribution of novel functional elements across MAF bins **d**. Distribution of exonic variants by functional elements. FS, frameshift; sSNV (synonymous SNVs); nsSNV (non-synonymous SNVs).

### Variant Prioritization and prediction of mutation burden

Potentially pathogenic SNVs were identified by selecting those that had at least 10 predictions of deleteriousness (**Table S4**). We also observed that 8 of the genes identified by at least 10 predictions in ANNOVAR harboured additional loss-of-function (LOF) variants according to snpEFF (**Table S4**). A trimmed list of five SNVs that were further classified as “damaging” by FATHHM is hereby presented. The most deleterious mutations were found within the *ACTRT2, HOXD12, ABCB5, ATP8B4* and *ABCC12* genes (**Table 1**).

**Table 1.** The most deleterious nonsynonymous single nucleotide variants.

| CHR | Position | ID | cDNA change | AA change | Gene | CADD | Botswana | 1KGP | gnomAD_AFR |
| --- | --- | --- | --- | --- | --- | --- | --- | --- | --- |
| 1 | 3022425 | rs3795263 | exon1:c.G739A | p.G247R | *ACTRT2* | 16.1 | A=0.0013 | A=0.12 | A=0.044 |
| 2 | 176100737 | rs200302685 | exon2:c.G790C | p.E264Q | *HOXD12* | 34 | C=0.032 | . | C=0.00038 |
| 7 | 20727068 | rs111647033 | exon13:c.G1319C | p.R440P | *ABCB5* | 20.1 | C=0.026 | C=0.0004 | C=0.0022 |
| 15 | 49972713 | rs77004004 | exon13:c.C1112A | p.P371H | *ATP8B4* | 14.2 | T=0.019 | T=0.0038 | T=0.013 |
| 16 | 48139198 | rs113496237 | exon5:c.G796C | p.G266R | *ABCC12* | 19.2 | G=0.013 | . | G=0.000071 |

CHR: chromosome. AA change: amino acid change. 1KGP: The 1000 Genomes Project MAF. gnomAD\_AFR: MAF of an African population from the gnomAD database.

### Distribution of pathogenic SNVs in known HIV-1 specific host genes

Discrepancies in pathogenic SNV proportions were observed between HIV-1 positive (HIV-1 cases) and HIV-1 negative (HIV-1 controls) in the Sec1 Family Domain Containing 1 (*SCFD1*), Histone Cluster 1 H4 Family Member B (*HIST1H4B*), Histone Cluster 1 H4 Family Member A (*HIST1H4A*), Immunoglobin Superfamily Member 21 (*IGSF21*), Nuclear Cap Binding Protein Subunit 2 (*NCBP2*) and Zinc Finger DHHC-Type Palmitoyltransferase 19 (*ZDHHC19*) genes. Lower proportions were observed for *SCFD1, HIST1H4B, HIST1H4A* and *ZDHHC19* genes, and higher proportions were observed for *IGSF21* and *NCBP2* genes in HIV-1 cases (**Figure 2**).

**Figure 2.**
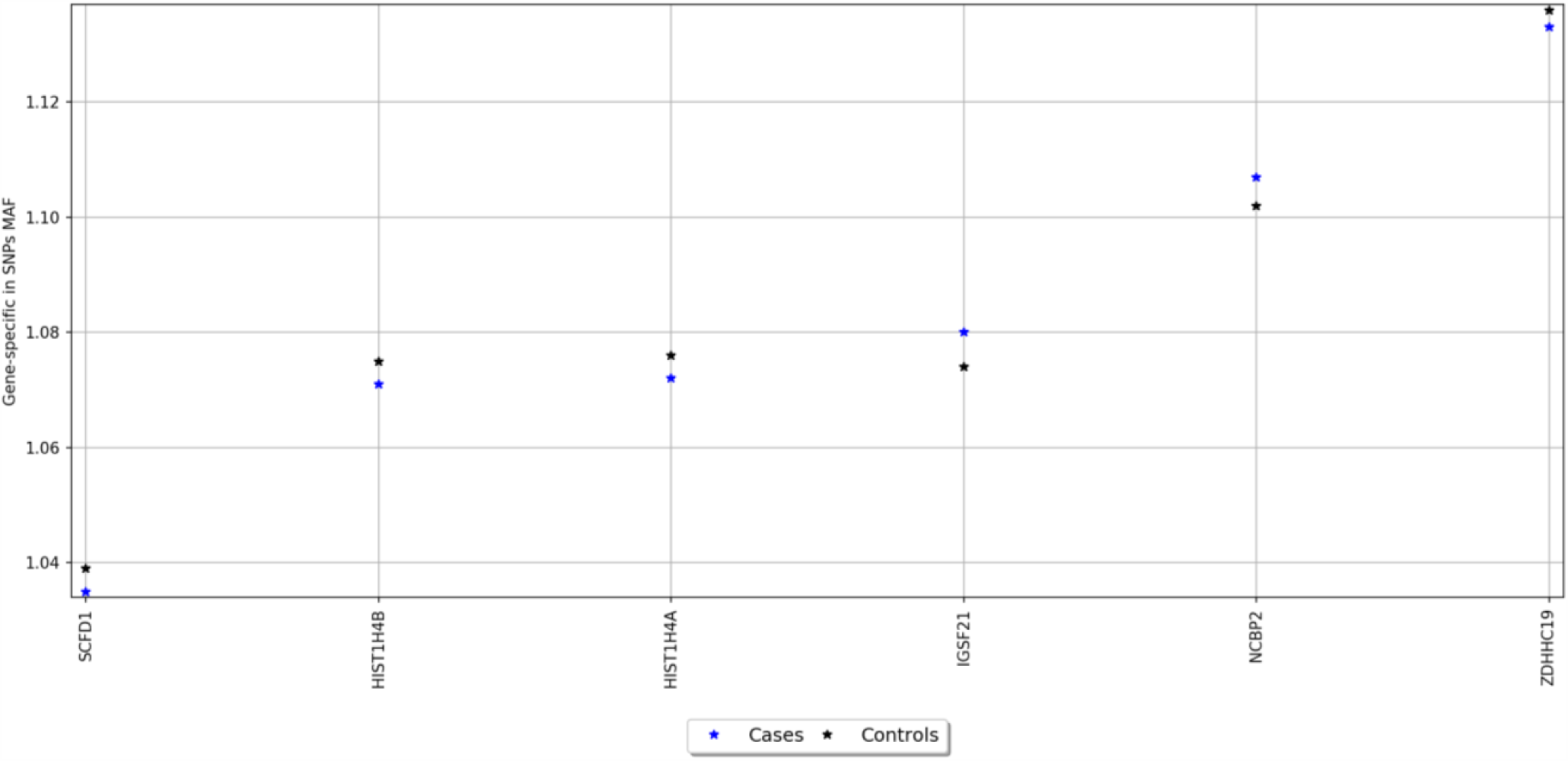
Distribution of pathogenic SNVs in known HIV-1 specific host genes.

### Pathways enrichment analysis and gene-gene interactions

The 24 genes harbouring the potentially pathogenic variants were subjected to enrichment analysis using GeneMANIA (39) and Enrichr (40) bioinformatics tools to identify biological processes and pathways putatively affected (Figure 8, Table 4). To successfully enrich for biological processes and pathways, the identified genes were used to “fish” 20 more related genes that are predicted to physically interact, co-express and co-localize with the identified genes (Figure 8).

The products of the identified genes were predicted to perform the following biological processes: gluconeogenesis, hexose and acyl-COA biosynthesis (**Table 4**). These gene products are localized within the oxoglutarate dehydrogenase complex and the mitochondria. The predicted molecular functions of these gene products were catalysis of peptidase, hydro-lyase, alcohol dehydrogenase and ATPase activities. The affected pathways included the glycolysis and gluconeogenesis, Krebs cycle, renal carcinoma, hypoxia-inducible factor 1 (HIF-1) signalling and folate biosynthesis pathways (**Table 2**). The identified genes were found to be associated with Pyruvate dehydrogenase complex deficiency (PDCD). One of the identified genes tumor susceptibility 101 (*TSG101*) was also found to be associated with human immunodeficiency virus 1 (HIV-1), albeit not statistically significant (**Table 2**).

**Table 2.**
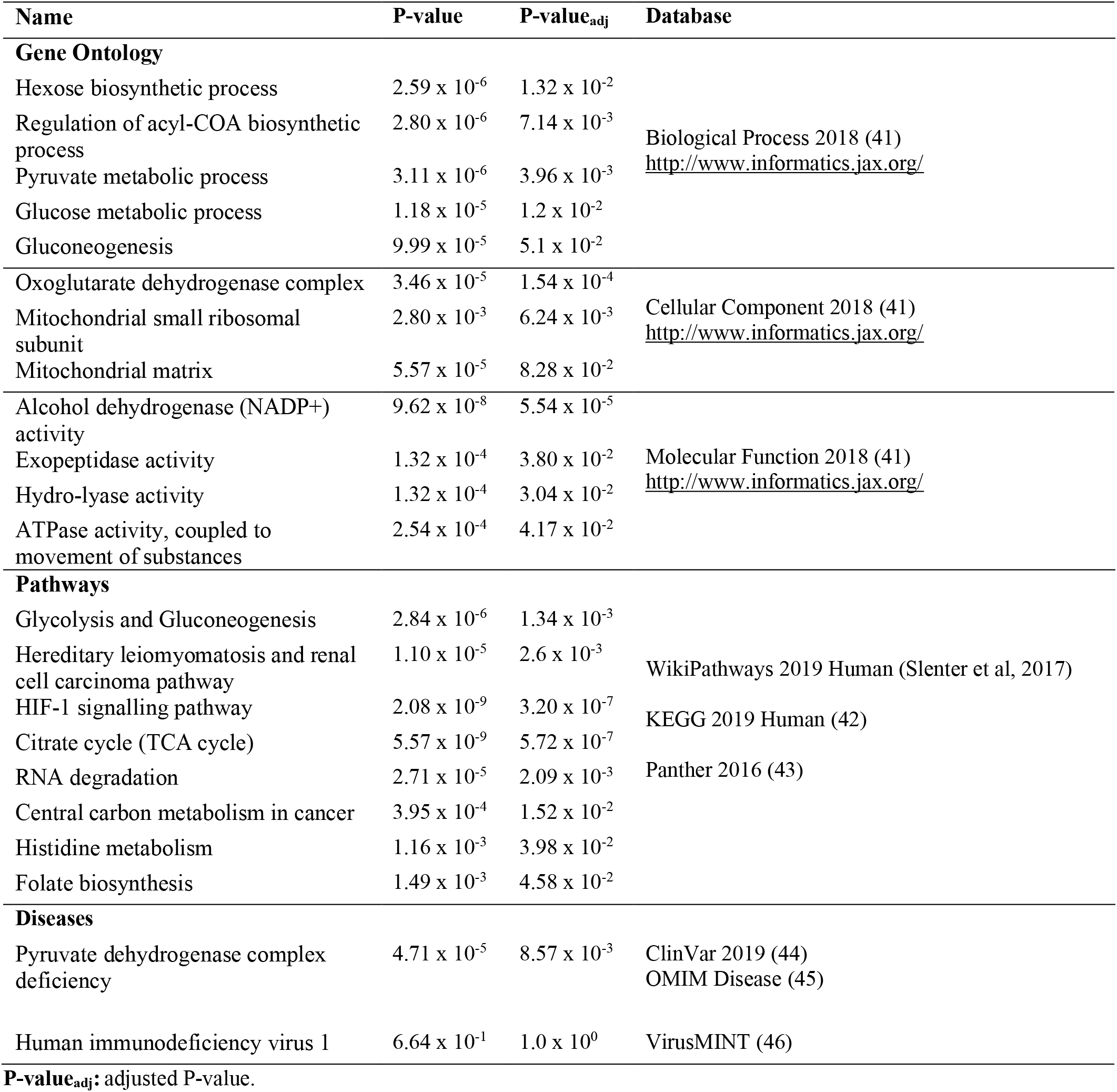
Enrichr gene-set enrichment of the genes harbouring the prioritized mutations.

### Population diversity

#### Principal components analysis (PCA) and admixture analysis

Population sub-structure was not observed within the Botswana study population. The plots of the first 3 PCs show a homogeneous mix of individuals from the HIV positive and the HIV negative groups with 3 outliers (Figure 4).

**Figure 3.**
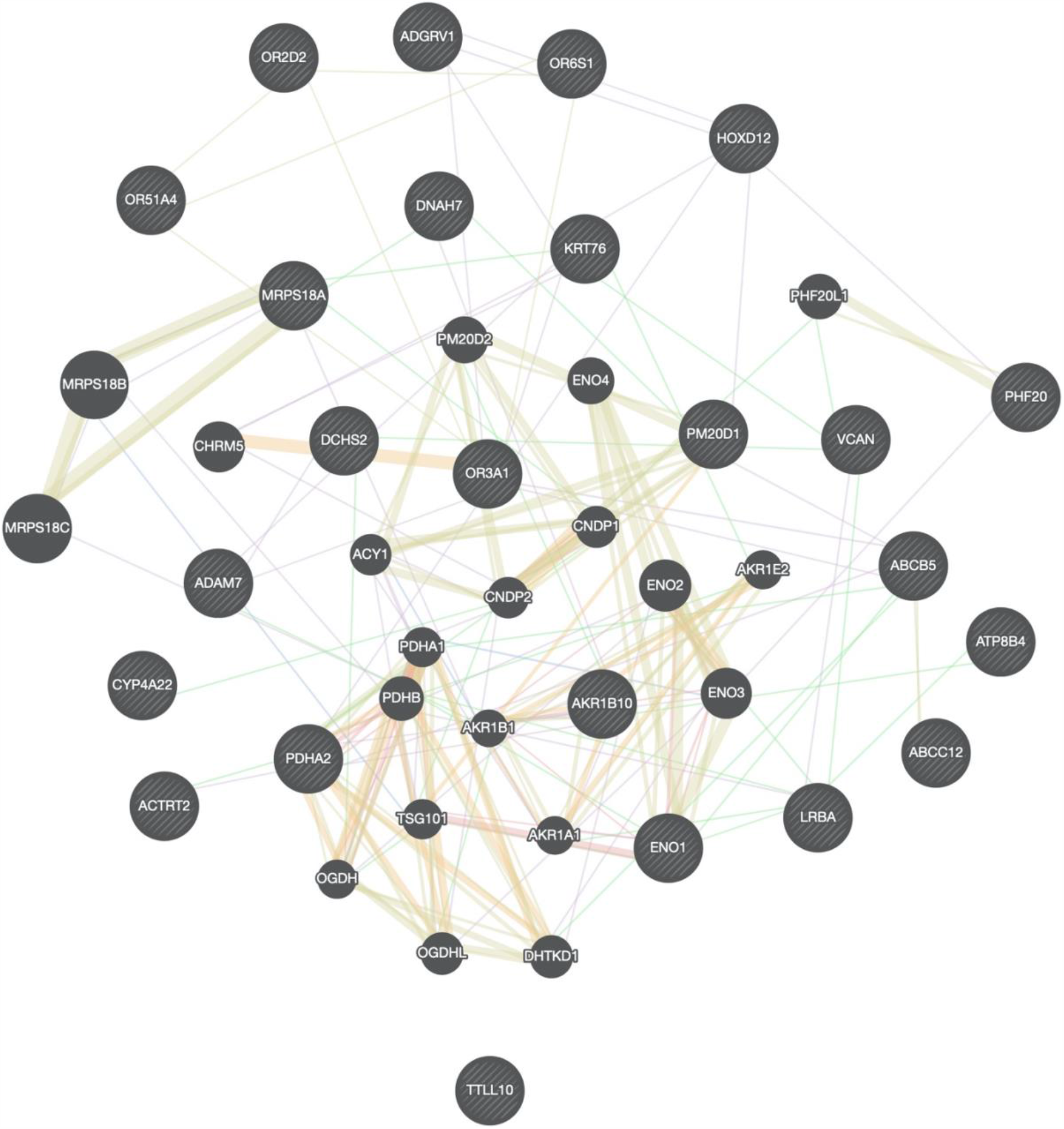
Gene-gene interaction network of genes harbouring the most deleterious variants. The different colours of branches represent how the genes are related; pink: physical interactions, purple: co-expression, orange: predicted, navy blue: co-localization, blue: Pathway, green: Genetic interactions, yellow: shared protein domains. Black and stripped nodes: genes provided as input into GeneMANIA. Black only nodes: genes predicted by GeneMANIA to interact with the input list. Connecting lines represent interactions.

**Figure 4.**
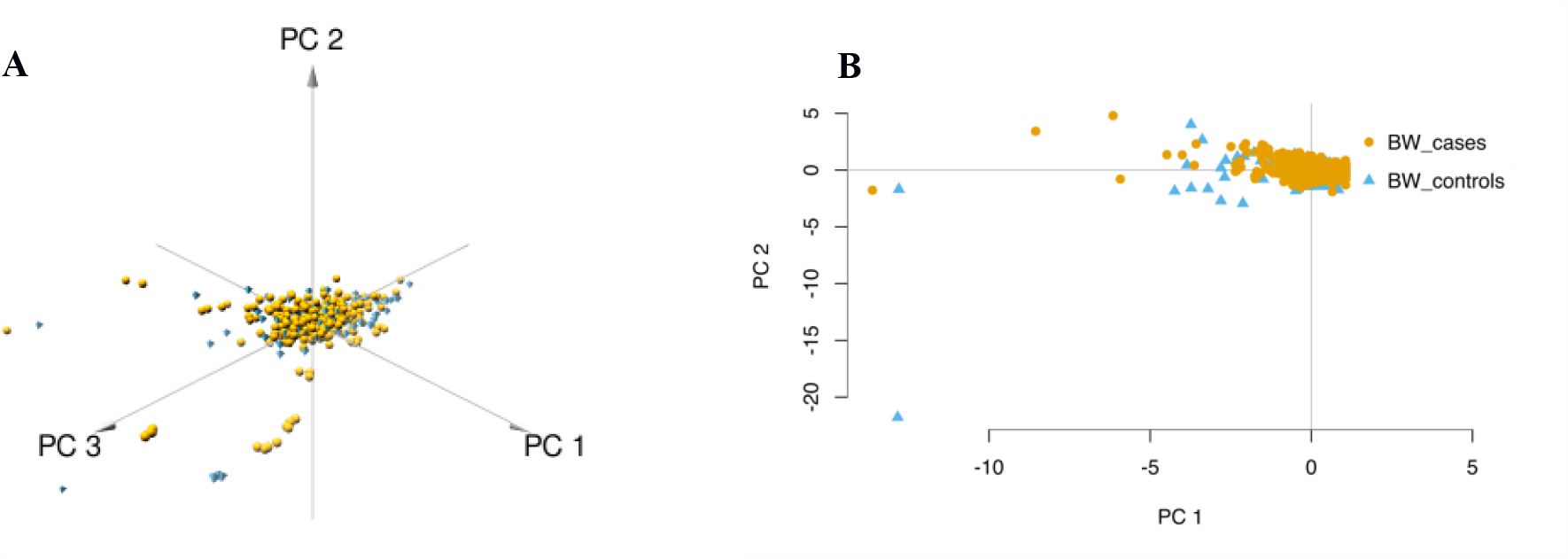
Principal component plot depicting population substructure of HIV-1 positive/negative individuals from Botswana. A depiction of population substructure of Botswana with a 3D plot PCs 1,2 and 3 (A) and 2D plot of PC2 against PC1 (B) showing cases (HIV-1 positive) in bright brown and controls (HIV-1 negative in blue).

The Botswana population formed a cluster with other African populations of the Niger-Congo ethnolinguistic phylum, away from the other ethnicities (**Figure 5**). We also assessed the genetic relationship between Batswana, other Niger-Congo populations and the Khoe-San. We see in **Figure 6** that Batswana and the Niger-Congo Bantu South formed a separate cluster from other Niger-Congo populations, with a dispersion towards the Khoe-San.

**Figure 5.**
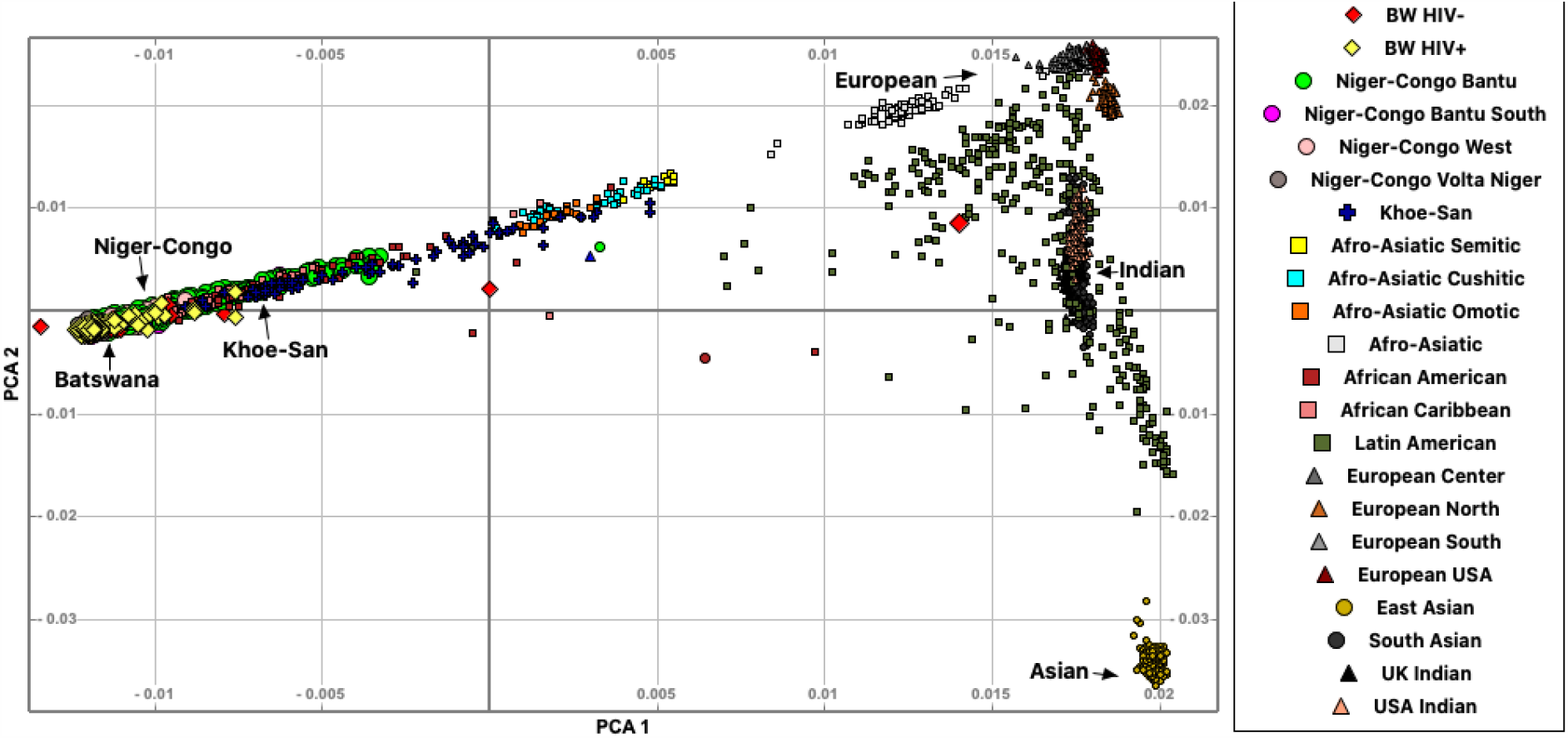
A PCA plot of the genetic relationship of the Botswana population with 20 world-wide ethnicities. The points on the PCA plot represent each individual. Botswana individuals (known as Batswana) are shown in diamond. Botswana HIV-1 negative (BW HIV-) individuals are shown with red diamonds, while Botswana HIV-1 positive (BW HIV+) individuals are shown with yellow diamonds.

**Figure 6.**
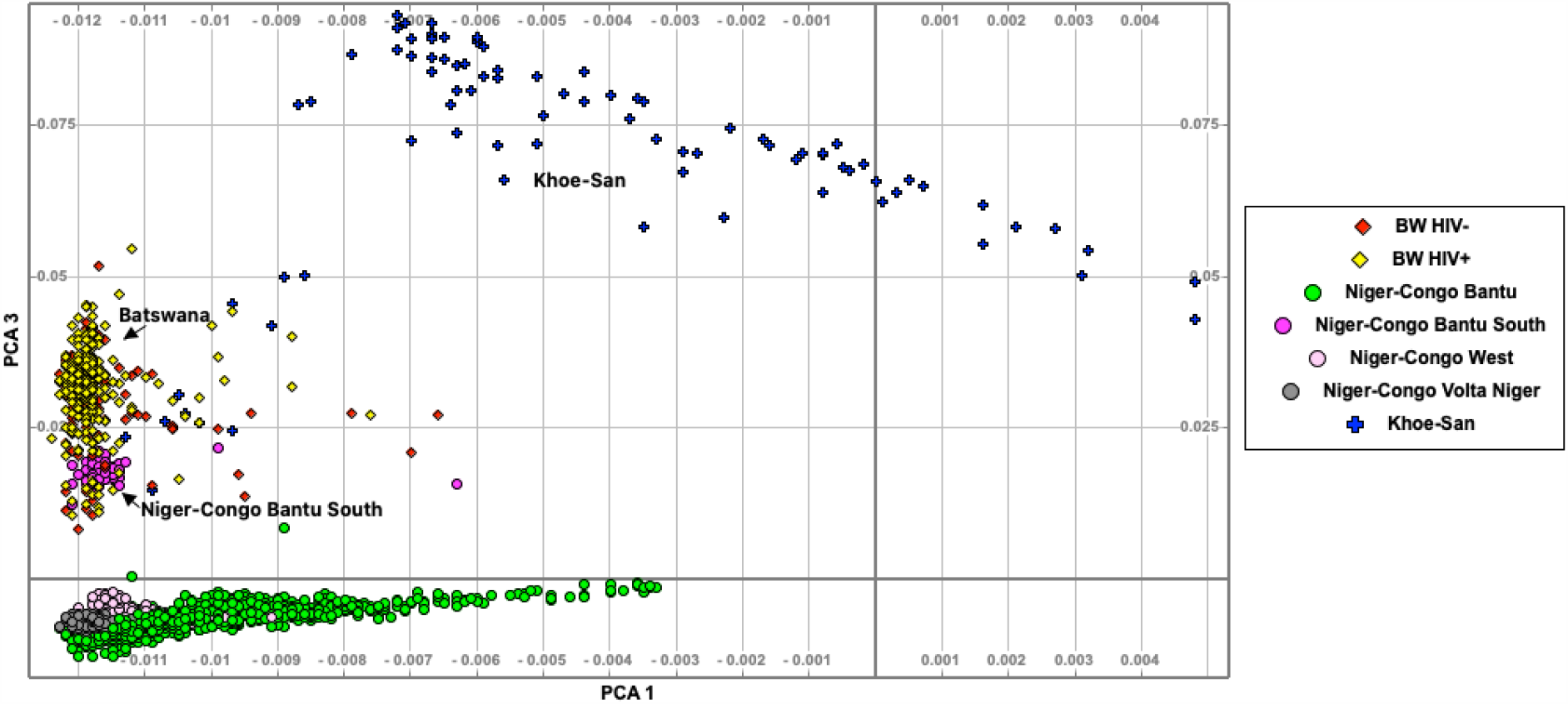
A PCA plot of the genetic relationship of Batswana, other Niger-Congo populations and the Khoe-San. Botswana samples are in the convex of Khoe-San and Bantu, confirming the genetic contribution of both Bantu and Khoe-San in Botswana.

Given the results in Figure 3, we performed admixture analysis to estimate the individual fraction of genetic ancestry. Batswana assessed in this study show admixture of the following ancestry proportions: Niger-Congo (65.9%), Khoe-San (32.9%) and Europeans (1.1%)

#### Population-based genetic distance (F_ST_)

The pairwise F_ST_ results accentuates what was observed in assessment of global population structure. The heatmap and hierarchical clustering shows two distinct clusters separating into the Eurasian and African clades. A sub-clade that branches into the Niger-Congo populations and the Khoe-San population was observed. An inner sub-clade that separates Southern Bantu-speakers (including the Botswana population) from other Niger-Congo population is also observed (Figure 8).

**Figure 7.**
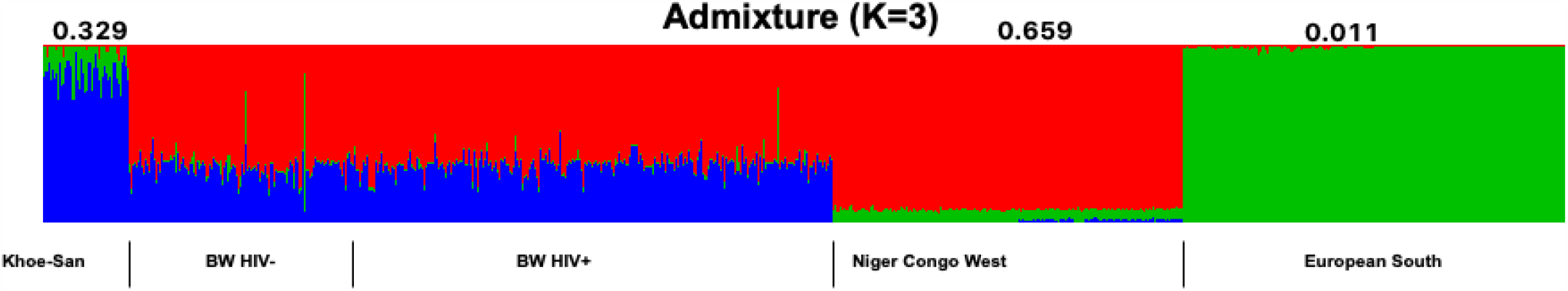
Genome-wide admixture proportions of Botswana. Khoe-San, Niger-Congo and European populations were used as proxy ancestral populations that may have potentially contributed to the genetic architecture of Botswana.

**Figure 8.**
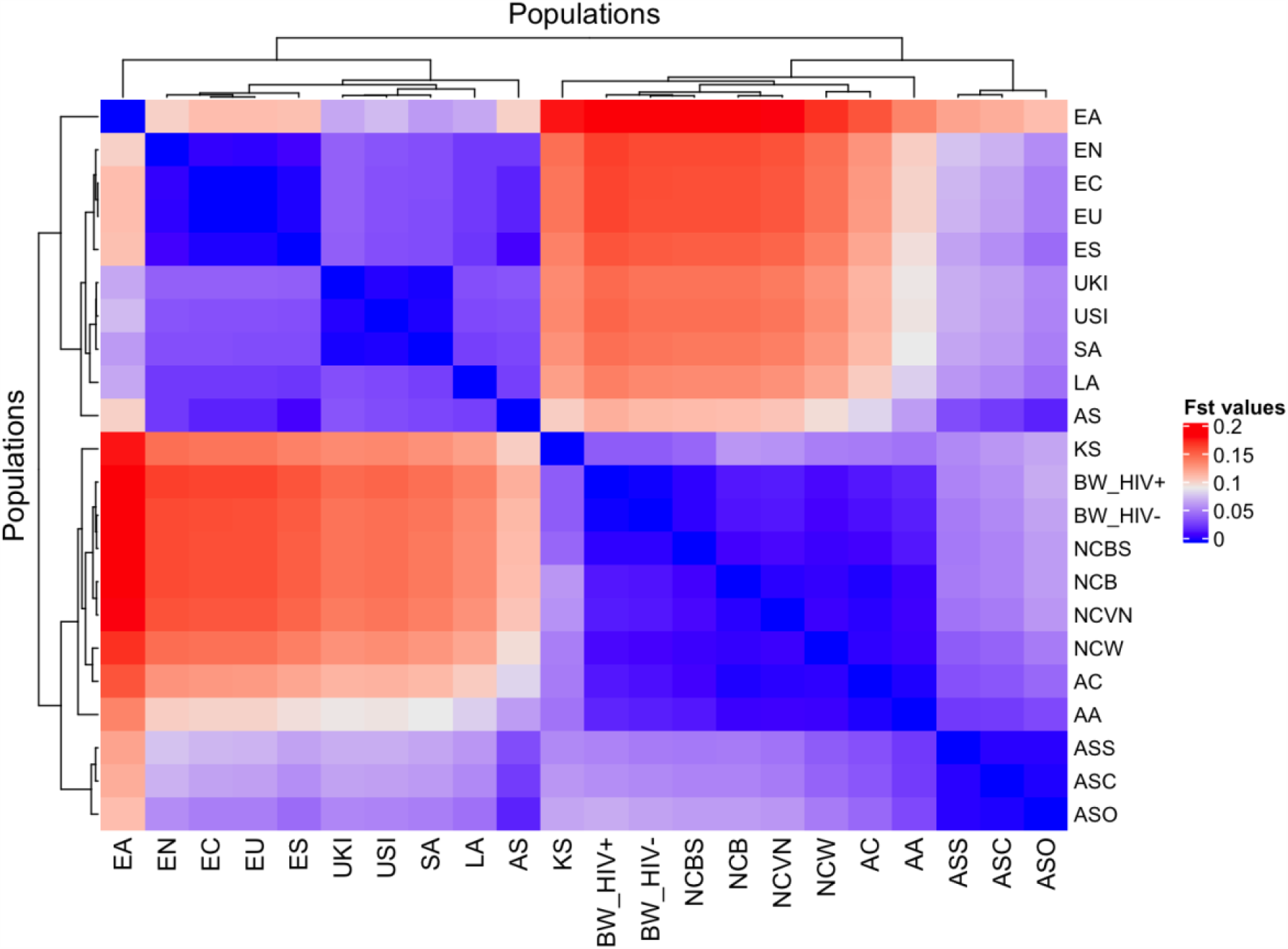
Pairwise genetic distance between the Botswana HIV-1 positive/negative population and 20 world-wide ethnicities. This is a heatmap and dendrogram of F_ST_ values showing pairwise genetic divergence between populations. The blue shade represents similarity while the red shade represents divergence between the populations. The populations are AA: African-American, AC: African-Caribbean, AS: Afro-Asiatic, ASC: Afro-Asiatic Cushitic, ASO: Afro-Asiatic Omotic, ASS: Afro-Asiatic Semitic, LA: Latin American, KS: Khoe-San, BW_HIV+: Botswana HIV-1 positive, BW_HIV-: Botswana HIV-1 negative, NCB: Niger-Congo Bantu, NCBS: Niger-Congo Bantu South, NCVN: Niger-Congo Volta Niger, NCW: Niger-Congo West, EN: European North, ES: European South, EU:USA European, EC: European center, EA: East Asian, SA: South Asian, UKI: UK Indian and USI: USA Indian.

### Genetic relatedness and runs of homozygosity

The IBD analysis revealed that none of the study participants were related. Our results further showed diversity in ROH segments among African populations, and between the African populations and non-African populations (Figure 9). Generally, the Niger-Congo populations (including the Botswana HIV-1 positive/HIV-1 negative cohort) had lower ROH lengths and less abundant ROH segments than the European, Asian, Indian, Latin-American and Khoe-San populations (Figure 9).

**Figure 9.**
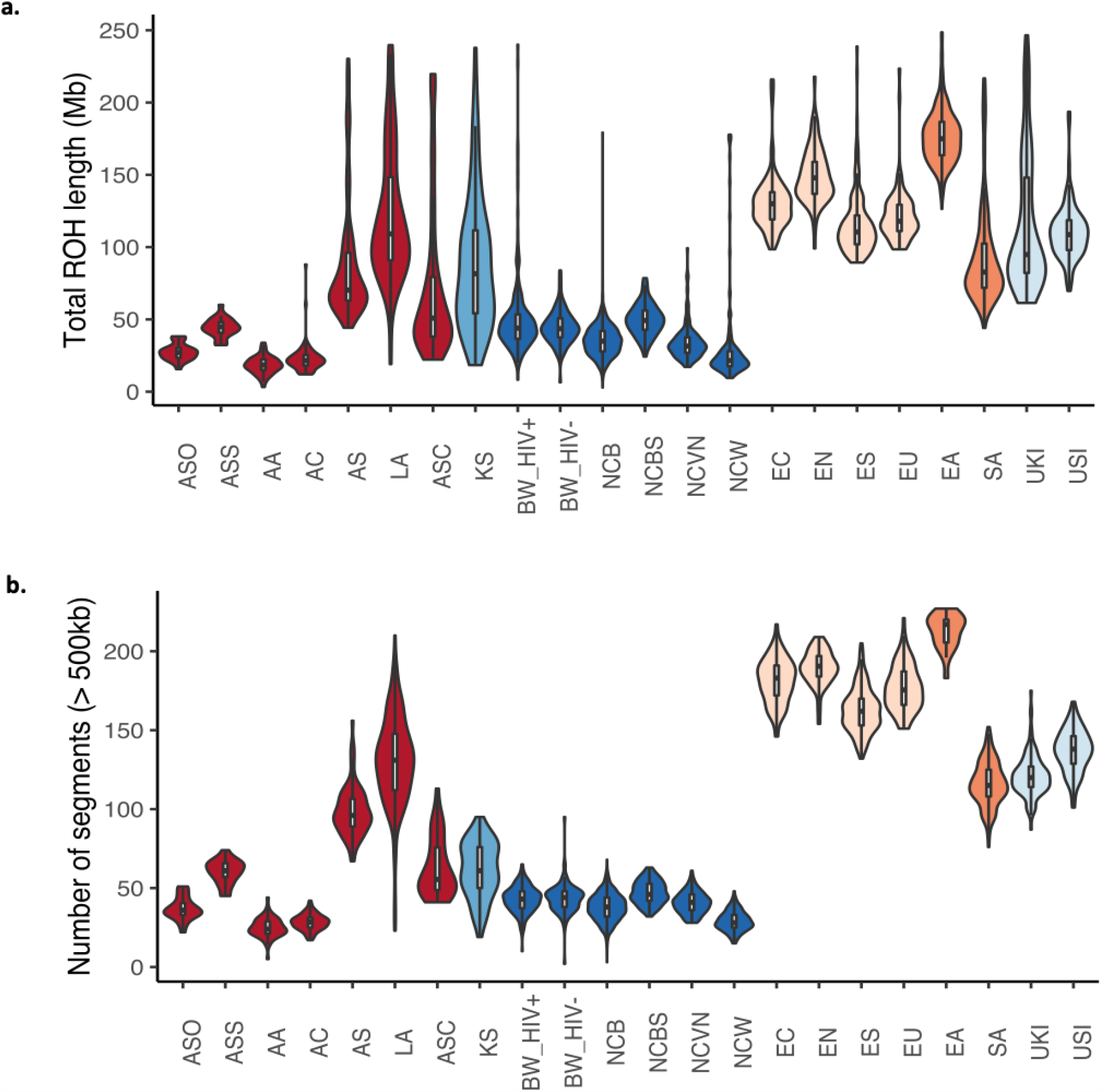
The lengths and number of runs of homozygosity (ROH) segments across different global ethnic groups. Violin plots showing median the lengths (in Mb) and number of ROH. The colours represent different super-groups: Mixed populations (African-American (AA), African-Caribbean (AC), Afro-Asiatic (AS), Afro-Asiatic Cushitic (ASC), Afro-Asiatic Omotic (ASO), Afro-Asiatic Semitic (ASS) and Latin American (LA)) in dark-red, Khoe-San (KS) in light blue, Niger-Congo in navy blue (Botswana HIV-1 positive (BW_HIV+), Botswana HIV-1 negative (BW_HIV-), Niger-Congo Bantu (NCB), Niger-Congo Bantu South (NCBS), Niger-Congo Volta Niger (NCVN) and Niger-Congo West (NCW)), Europeans (European North (EN), European South (ES), USA European (EU), European center (EC)) in light orange, Asians (East Asian: EA and South Asian: SA) in orange and Indians (UK Indian (UKI) and USA Indian (USI)) in very light blue.

#### Comparison of genome-wide admixture proportions between HIV-1 positive and HIV-1 negative groups

The genome-wide genetic proportions of Khoe-San ancestry in Botswana cases (HIV positive individuals) was significantly higher (0.336 ± 0.003 vs 0.315 ± 0.005, p-value = 0.002) than that observed in Botswana controls (HIV negative individuals) (Table 6). There was no significant difference in the genome-wide genetic proportions of the Niger-Congo and European ancestries when comparing Botswana cases to Botswana controls (Table 5).

## DISCUSSION AND CONCLUSION

Of the 27.7 million variants identified from 30X depth whole genomes of 390 individuals of Botswana. A critical and convenient QC metric to measure the quality and accuracy of genomic variation data is the TI/TV ratio (47). The average TI/TV ratio of this set of variants was 2.1. This TI/TV ratio is within the expected range for human whole genome data which is ∼2.0-2.1, meaning that the data is devoid of false positives. As observed previously (27), intergenic variants had the highest frequency, followed by intronic variants and non-coding RNA (ncRNA) variants. Thirteen percent (2,789,599) of the discovered SNVs were novel. This number of previously uncaptured genetic variation highlights a potential of identifying population-specific variations through WGS. Whole genome sequencing also offers an opportunity to identify intronic variants and variants within non-coding regions. To this effect 1,066,166 intronic and 178,178 (ncRNA) novel variants were identified.

Recent human population expansion has resulted in a skewness towards excessive rare variants. This means that rare variants constitute a large part of the human genomic variations (48–52). Hence it is not surprising that a majority of the novel variants identified in the current study were very rare, occurring only once in the dataset (**Table S3, Figure 1a** and **c**). A substantial number of the exonic variants were nonsynonymous, stop gain and stop loss variants (**Figure 1d** and **Table S2**). These three types of mutations respectively cause a change in the amino acid and lead to an abnormal truncation or elongation of the protein, all leading to a change in the conformation or function of the encoded protein (53). These changes have a potential to disturb normal biological processes and cause disease. In fact, a lot of genetic diseases are caused by nonsynonymous mutations.

Variants classified as damaging by at least 10 deleteriousness tools were further prioritized with FATHHM score (**Table 3**). Here 5 variants within 5 genes were predicted to be the most deleterious (rs3795263 in the *Actin Related Protein T2* (*ACTRT2*) gene, rs200302685 in *homeobox D12* (*HOXD12*) gene, rs111647033 in ATP binding cassette subfamily B member 5 (*ABCB5*) gene, rs77004004 in *ATPase phospholipid transporting 8B4* (*ATP8B4*) gene and rs113496237 in *ATP Binding Cassette Subfamily C Member 12* (*ABCC12*) gene. The product of *ACTRT2* gene may be involved cytoskeletal organization (54). The rs3795263 variant was previously identified as harmful and associated with a severe form of tick-borne encephalitis virus infection (55). The *HOXD12* gene belongs to the homeobox (*HOX*) family of genes that encode transcription factors involved in regulation of embryonic development (54,56). The exact role of HOXD12 is unknown (54). The *HOX* genes have been implicated in maintenance and control of HIV-1 latency through epigenetic regulation (57).

**Table 3.** Comparison of the mean genetic ancestry proportions of Botswana estimated with ADMIXTURE between HIV-1 positive and HIV-1 negative groups.

| Ancestry | Genome-wide ancestry contribution (mean ± SE) | | | Mean comparisons of cases vs controls (P-value) |
| --- | --- | --- | --- | --- |
|  | All samples | Cases (HIV+) | Controls (HIV-) |  |
| Khoe-San | $0.329 \pm 0.003$ | $0.336 \pm 0.003$ | $0.315 \pm 0.005$ | 0.002 |
| Niger-Congo | $0.659 \pm 0.003$ | $0.657 \pm 0.003$ | $0.665 \pm 0.007$ | 0.328 |
| European | $0.0114 \pm 0.003$ | $0.00727 \pm 0.002$ | $0.0202 \pm 0.007$ | 0.076 |

Estimation of the mean and SE (standard error of the mean) of ancestry proportions from each of the 3 populations contributing to the admixture of Botswana.

The *ABCB5* gene belongs to the ATP-binding cassette (ABC) family that encodes proteins responsible for transmembrane transport of molecules including drugs such as doxorubicin (54). *ABCB5* is thought to also mediate chemoresistance of doxorubicin in malignant melanoma, (58). The *ATP8B4* gene encodes an ATPase protein that is responsible for phospholipid translocation in the cell membrane (54). The *ABCC12* gene also encodes an ABC protein responsible for transmembrane transport of molecules. Overexpression of the *ABCC12* gene has been associated with breast cancer (54). Some members of the *ABC* family regulate the efflux of HIV-1 antiretrovirals from intracellular compartments (59,60). Biological pathways potentially affected by the products of these putatively deleterious genes and their interactome are discussed in subsequent paragraphs.

The minor allele frequencies of the *HOXD12* rs200302685, *ABCB5* rs111647033, *ATP8B4* rs77004004 and *ABCC12* rs113496237 variants in the Botswana data were generally higher when comparing to the gnomAD and the 1000 Genomes Project data. While the MAF for the *ACTRT2* rs3795263 variant was lower than in the gnomAD and the 1000 Genomes Project data. This highlights that MAFs do vary per ethnicity which could affect the risk of disease differently between populations (**Table 1**).

Gene-set enrichment and functional analysis revealed the following pathways that were enriched for with the putatively deleterious genes: glycolysis and gluconeogenesis, Krebs cycle, renal carcinoma, Hypoxia-inducible factor 1 (HIF-1) signalling, RNA degradation, Histidine metabolism and folate biosynthesis pathways (**Table 2**). The *pyruvate dehydrogenase* (*PDH*), *enolase* (*ENO*) and *aldo-keto reductase* (*AKR1*) genes (**Figure 8, Table 2**) were significantly associated with glycolysis and gluconeogenesis (1.34 x 10^−3^).

Both glycolysis and gluconeogenesis are glucose metabolism pathways; glycolysis is the catabolism of glucose (or glycogen) into pyruvate, while gluconeogenesis is the anabolism of pyruvate (from mainly proteins) into glucose (61,62). The PDH genes were also significantly associated with the Kreb’s (Tricarboxylic Acid - TCA or Citric Acid) Cycle (p = 5.72 x 10^−7^). Glycolysis, gluconeogenesis and the TCA cycle are involved in energy (mostly in the form of Adenosine triphosphate - ATP) production, carried out in the cytoplasm or mitochondria of eukaryotes (**Table 2**) (61,62). In cells where there is low oxygen (a condition known as hypoxia), HIF-1 gets activated and triggers energy production through anaerobic glycolysis (63).

Glycolysis and TCA intermediates are used as precursors for macromolecule synthesis in hypoxic conditions such as cancer (64,65). This may explain the association of the *PDH* genes with HIF-1 signalling and cancer pathways (**Table 2**). Amino acids are also made from intermediates of TCA, glycolysis and the pentose phosphate pathways (66). This may explain the significant association of histidine biosynthesis with genes that are in connection with glycolysis and TCA pathways (**Figure 8, Table 2**).

The association of *AKR1* genes (**Figure 8**) with alcohol dehydrogenase (NADP+) activity and folate biosynthesis (**Table 2**) could be explained by that the alcohol dehydrogenases catalyse the reduction of NADP+ to NADPH (67). This reaction also takes place within glycolysis, gluconeogenesis and pentose phosphate pathways (61,66). Furthermore, there is also evidence of NADPH being produced from folate metabolism (68). The human polynucleotide phosphorylase (hPNPase^old-35^) is an evolutionary conserved RNA-degradation enzyme that has homologues in organisms such as *Escherichia coli* and yeast (69,70). In *E. coli* PNPase forms part of the degradosome with enolase and a helicase (71). This link between enolase and the evolutionary conserved PNPase may explain the association of the *ENO* genes with RNA degradation (**Table 2**). The degradation of HIV-1 mRNA in HIV-1 infected cells is important in suppressing HIV-1 replication (72). Moreover, ENO-1 has been shown to prevent HIV-1 reverse transcription and ultimately decrease HIV-1 infectivity (73).

Lower proportions of potentially pathogenic SNVs were observed for *SCFD1, HIST1H4B, HIST1H4A* and *ZDHHC19* genes, and higher proportions were observed for *IGSF21* and *NCBP2* genes in HIV-1 cases (**Figure 7**). The *IGSF21* gene encodes a cell receptor that is a member of the immunoglobulin superfamily (54). An intron variant rs2883821 within the *IGSF21* gene (chromosome 1) was reported to be associated with tenofovir pharmacokinetics and increased HIV-1 viral load (74). The *NCBP2* gene encodes a protein that is part of the nuclear cap-binding protein complex (CBC). The CBC binds to the pre-mRNA and is involved in various processes such as splicing, transcription and nonsense-mediated mRNA decay (54). A splice site variant rs548853 within the *NCBP2* gene (chromosome 3) has been associated with decrease in viral load (74). Since the proportion of pathogenic variants within the *IGSF21* and *NCBP2* genes in HIV-1 cases is higher than in HIV-1 control, this corroborates with the previous study, that *IGSF21* gene may harbour risk alleles.

The product of the *SCFD1* gene (chromosome 14) plays a role in SNARE-pin assembly and transport of molecules from the Golgi apparatus to the endoplasmic reticulum (54). SCFD1 interacts with other Golgi proteins and possibly affects HIV-1 replication through regulation of glycosylation. A reduction in the level of SCFD1 was observed to reduce HIV-1 infection (cell entry) (75). The *HIST1H4A* and *HIST1H4B* genes (chromosome 6) encode histones, these are proteins that bind to DNA and assist in compacting it into nucleosomes which are the basic repeating units of a chromatin. Therefore histones play critical roles in organizing chromatin structure and gene expression (54,76,77). HIV-1 induced a modulation of the chromatin signalling network that involved HIST1H4A through epigenetic modifications (78).

The *ZDHHC19* gene (chromosome 3) encodes a palmitoyl acyltransferase, an enzyme that mediates palmitoylation of signal transducer and activator of transcription 3 (STAT3) (54). Palmitoylation is a lipid modification process in which a fatty acids such as palmitate are attached to a cysteine residue of a protein to regulate its attachment and localization to the cytoplasmic membrane (79–81). In HIV-1 infection, STAT3 has been found to promote inflammation (82) and also to promote antiviral immune responses (83,84). According to the GWAS Catalog (74) a variant rs11924930 within the *ZDHHC19* gene has been associated with HIV-1 susceptibility in a GWAS study of a Malawi population (85). However, the effect of this variant is not publicly available. In the current study, lower proportions of pathogenic SNVs were observed for the *SCFD1, HIST1H4B, HIST1H4A* and *ZDHHC19* genes in HIV-positive individuals. This might indicate that the minor alleles are protective against HIV-1 infection.

The assessment of population substructure through PCA revealed no evidence of substructure in the study population of Botswana (**Figure 4**). The study participants were recruited from three districts in the southern part (Southern, Kweneng and South-East) of Botswana. Although the sampling site does not span the whole of Botswana, the current findings have positive implications for genetic epidemiology in the southern part of Botswana.

Population substructure can mask true genetic associations and also lead to false discovery of causal (or modifier) variants (14–17). To find minimal or no substructure in the study populations will minimize false positives in subsequent genetic association analyses. Furthermore, population-specific interventions against HIV-1 can be employed for this part of Botswana which will minimize costs that may arise in personalized medicine.

The PCA (**Figure 5**) and F_ST_ values (**Figure 8**) show that there was a clear distinction between African populations and European populations. The population of Botswana clustered with other Niger-Congo populations and showed a dispersion towards the Khoe-San population (**Figure 6**). The Botswana population showed a closer affinity with the Niger-Congo Bantu South (Zulu) population. This is expected as a close affinity of the Sotho with the Niger-Congo Bantu South (Zulu) has previously been reported (27). Batswana are members of the Sotho-Tswana clan of Southern Africa that includes the Sotho (of Lesotho and South Africa) and Batswana (of Botswana and South Africa) (34,35).

Major events such as the “Bantu expansion” and Eurasian migration into Southern Africa have shaped the genetic landscape of the region. These events have led to varying degrees of admixture of the migrant groups and indigenous population (14,25–31). These previous findings are congruent with the current study that reports a 3-way admixture of Niger-Congo (65.9%), Khoe-San (32.9%) and European (1.1%) populations observed in the Botswana population (**Figure 7** and **Table 3**).

We found no evidence of consanguinity in the Botswana HIV-1 positive/HIV-1 negative cohort as defined by less abundant segments and lower lengths of ROH in comparison to non-African populations and the Khoe-San (**Figure 9**). This finding is supported by the previous observation of no extended ROH lengths in a Botswana HIV positive cohort (86). Among the Niger-Congo populations, the median ROH length in the Botswana HIV-1 positive/HIV-1 negative and the Niger-Congo Bantu South were significantly higher (p-value = 2.2 x 10^−16^) than of the Niger-Congo Bantu, Niger-Congo West and the Niger-Congo Volta Niger (**Figure 9, Table S6**). These results are consistent with what was observed by Choudhury *et. al*., who observed that the Niger-Congo Bantu population of Southern Africa had the highest lengths of ROH compared to Niger-Congo populations of East, Central West and West Africa (27).

The mean proportion of Khoe-San ancestry was higher in HIV-1 cases than in controls (**Table 3**). Host genetics studies have previously linked Khoe-San ancestry to susceptibility to tuberculosis (TB) (87). Could this be yet another association of the Khoe-San ancestry with an infectious disease as with TB? If the answer is an affirmation then from a public health standpoint, this could mean that efforts of HIV-1 prevention should be elevated among people of Khoe-San ancestry. However, this conclusion seems to be implausible since when we regressed HIV-1 status against the ancestry proportions none of the ancestries showed association with HIV-1 status.

It may be concluded that no substructure was observed in Botswana. This is good because if present, population structure cannot be fully corrected in genetic association studies. Admixture of Niger-Congo, Khoe-San and European populations was observed in the Botswana population. This is not surprising as Botswana is one of the countries with the largest number of the Khoe-San. The Khoe-San are known to be the indigenous people of Southern Africa. Overtime the Khoe-San are expected to have mingled and interbred with the Niger-Congo people of Botswana. Hence, this work shows the pivotal role played by genetics in the reconstruction of population histories. A limitation of this study is that the Botswana population had no ethnolinguistic labels, as such ethnicity inferences cannot be drawn from this study. Nevertheless, population structure and admixture could still be assessed as the algorithms used in this study are unsupervised machine learning methods and therefore can still give meaningful results.

This study is the first to use deep sequencing in efforts to delineate a complete genome map the human population of Botswana and evaluate the burden of human genomic mutations in Botswana. To this effect we identified single nucleotide variants which could potentially disrupt the function of 24 genes, the most deleterious (damaging) variants being *ACTRT2* rs3795263, *HOXD12* rs200302685, *ABCB5* rs111647033, *ATP8B4* rs77004004 and *ABCC12* rs113496237. Rare and low-frequency variants constituted the bulk of novel variants that were identified in this study. This was made possible by the unique potential of deep sequencing that offers an opportunity to discover rare variants. This is important because unlike Mendelian conditions, complex traits are influenced by many small-effect variants from different genetic loci, a concept known as polygenicity (88). The cumulative effect of rare variants plays an important role in the expression of complex traits such as HIV-1.

Glycolysis, TCA and hexo-pentose pathways emerged to be the most affected by the putatively deleterious variants. These are critical physiological pathways responsible for energy production, amino-acid biosynthesis, immunity and tumorigenesis among other roles. There were disparities in proportions of pathogenic variants within previously HIV-1 associated genes: *SCFD1, HIST1H4B, HIST1H4A, ZDHHC19, IGSF21* and *NCBP2* in HIV-1 infected versus uninfected individuals. Of interest in these genes is the *ZDHHC19* gene that encodes palmitoyl acyltransferase involved in pathways such as viral immunity. The *ZDHHC19* gene was previously identified in a GWAS of HIV-1 susceptibility in Malawi. Though the effect of this gene in the previous study is known, the identification of the gene in the current study confirms that the gene may have an implication in the genetics of HIV-1 in African populations. The current study also suggests a substantial level of pleotropic effects in the genome of Batswana. Although the candidate genes have been linked to HIV-1 infection, the same genes may also confer risk towards other health complications. This implies that the results may give insights into the potential interplay of genetic co-morbidities in the population of Botswana.

## MATERIALS AND METHODS

### Ethical approval

This study is part of a bigger protocol titled “Host Genetics of HIV-1 Subtype C Infection, Progression and Treatment in Africa/GWAS on determinants of HIV-1 Subtype C Infection” conducted by Botswana Harvard AIDS Institute Partnership. Ethics approval was obtained according to The Declaration of Helsinki. All participants consented to participate in the study. Institutional Review Board (IRB) approval was obtained for these samples from Botswana Ministry of Health and Wellness - Health Research Development Committee (HRDC) & Harvard School of Public Health IRB (reference number: HPDME 13/18/1) and the University of Cape Town - Human Research Ethics Committee (HREC reference number: 316/2019).

### Selection of study participants

This is a retrospective study that used samples from previous studies conducted at Botswana Harvard AIDS Institute Partnership between 2001 and 2007. Of the 390 participants, 265 were HIV-1 positive and 125 were HIV-1 negative. The participants were recruited from four locations within the southern region of Botswana (Mochudi, Molepolole, Lobatse and Gaborone) (Figure 10). The HIV-1 positive participants were previously part of the Mashi study (89,90), while HIV-1 negative participants were previously part of the Tshedimoso study (91).

**Figure 10.**
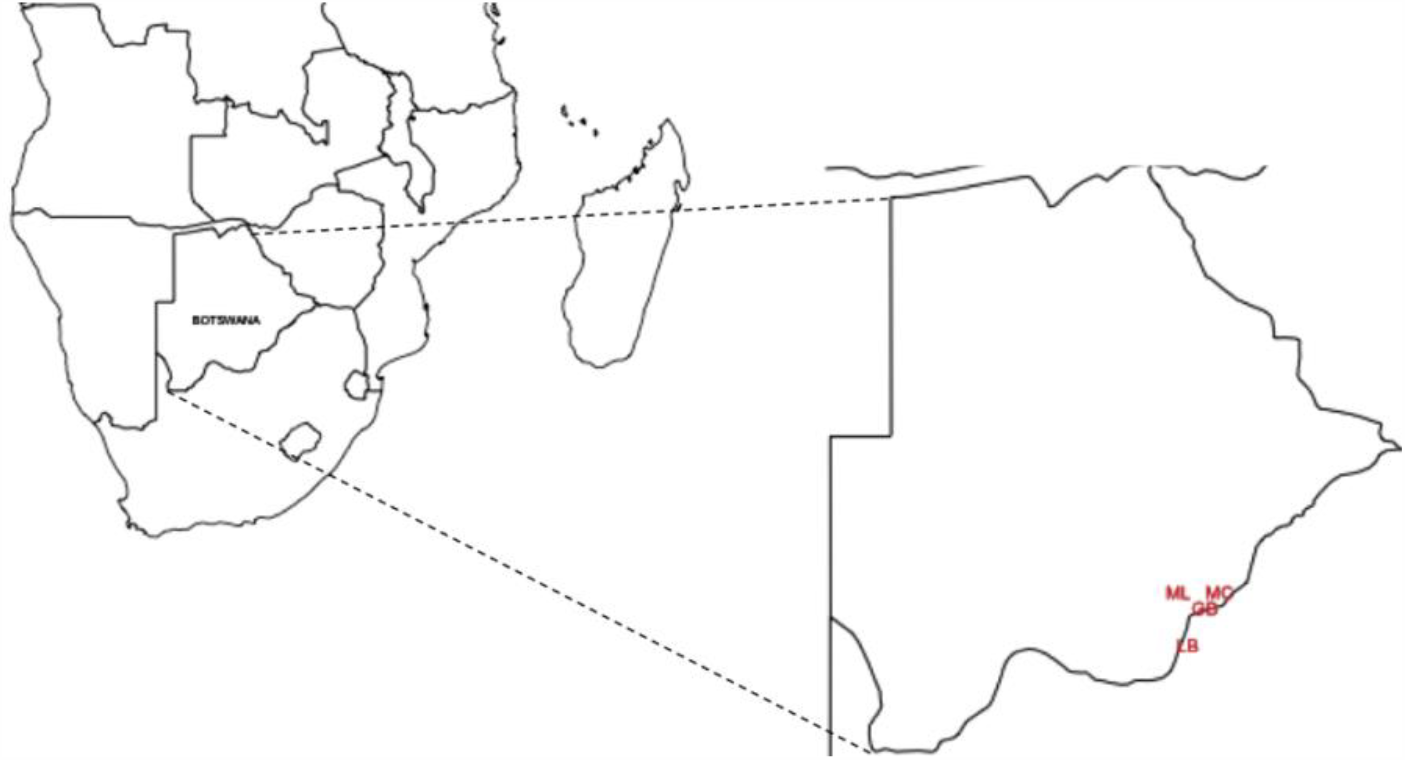
Whole genome sequencing sampling sites in Botswana. Botswana is located at the center of Southern Africa. The sampling sites are GB: Gaborone, LB: Lobatse, MC: Mochudi, ML: Molepolole. The map was produced with Maps package in R (92).

### DNA and Genomic characterisation

DNA was extracted from buffy coat using Qiagen DNA isolation kit following manufacturer’s instructions. DNA concentration was quantified using Nanodrop^®^ 1000 (Thermo Scientific, USA). Whole genome sequences of 394 Botswana nationals were generated using paired end libraries on Illumina HiSeq 2000 sequencer at BGI (Cambridge, MA, US).

### Variant Calling and Downstream Data Description

Quality assessment was performed on paired-end WGS (minimum of 30X depth) in FASTQ format (93) using FastQC (94). Low-quality sequence bases and adapters were trimmed using Trimmomatic with default parameters (95). The sequencing reads were aligned to the GRCh38 human reference genome using Burrows-Wheeler Aligner (BWA-MEM) (96,97) and post-alignment quality control including adding of read groups, marking duplicates, fix mating and recalibration of base quality scores was performed using Picard tools, SAMtools (98) and Genome Analysis Toolkit (99). Four samples (HIV-1 positive females) were excluded due to poor quality of sequences, the remaining dataset had 390 individuals. We have run FastQC on all final BAM files prior the variant calling, then we aggregated the results from FastQC into a single report by using MultiQC (100). All the sequences passed quality control.

We performed population joint calling (101,102) using two different population joint calling methods to leverage the quality and accuracy of our results: GATK HaplotypeCaller (47,99) and BCFtools (98). The variant call format (VCF) dataset was filtered using VCFTOOLS (103), GATK Variant Quality Score Recalibration and BCFtools. The specific filtering parameters employed for both call-sets have been detailed (**Supplementary information**). Downstream analyses were performed with GATK call-set and BCFtools call-set used as a validation set.

### Variants Annotation and Mutation Prioritization

The high confidence variants obtained from variant quality control, ANNOVAR (104) and snpEFF version 4.3T (105) were used to perform functional annotation in the Botswana HIV-1 positive/negative VCF file to determine whether SNPs cause protein coding changes and produce a list of the amino acids that are affected. We used ANNOVAR “2016Dec18” setting, where the population frequency, pathogenicity for each variant was obtained from 1000 Genomes exome (2), Exome Aggregation Consortium (106) (ExAC), targeted exon datasets and COSMIC (107). Gene functions were obtained from RefGene (108) and different functional predictions were obtained from ANNOVAR’s library, which contains up to 21 different functional scores including SIFT (109,110), LRT (111), MutationTaster (112), MutationAssessor (113), FATHMM (114,115), fathmm-MKL (114,115), RadialSVM (12), LR (12), PROVEAN (12), MetaSVM (12), MetaLR (12), CADD (12,116), GERP++ (117), DANN (104), M-CAP (104), Eigen (104), GenoCanyon (104), Polyphen2 HVAR (118), Polyphen2 HDIV (118), PhyloP (119) and SiPhy (119). From each resulting functional annotated data set, we independently filtered for predicted functional status (of which each predicted functional status is of “deleterious” (D), “probably damaging” (D), “disease_causing_automatic” (A) or “disease_causing” (D)) from SIFT, LRT, MutationTaster, MutationAssessor, FATHMM, fathmm-MKL, RadialSVM, LR, PROVEAN, MetaSVM, MetaLR, CADD, GERP++, DANN, M-CAP, Eigen, GenoCanyon, Polyphen2 HVAR, Polyphen2 HDIV, PhyloP, and SiPhy.

We prioritized the variants by retaining a variant only if it had at least 10 predicted functional status “D” or “A” out of 21 (120). To refine the results from this strategy, we re-applied FATHMM (114,115,121), a disease-specific weighting scheme, which uses a Hidden Markov Models prediction algorithm capable of discriminating between disease-causing mutations and neutral polymorphisms. FATHMM has been found to have the most discriminative power among other individual *in silico* mutation prediction tools (121). We identified further deleterious variants within the prioritized genes with snpEFF loss-of-function (LOF) module (105).

### Distribution of pathogenic SNVs in known HIV-1 specific host genes

Following the functional annotation of the discovered variants, we evaluated the share of pathogenic SNVs between HIV-1 positive and HIV-1 negative individuals from Botswana. We further classified the SNVs as pathogenic or population specific if their MAFs were lower than 5%. The proportion of pathogenic SNVs within a gene was defined as the count of observed pathogenic variants over the total number of variants in the given gene (120). We obtained a list of 730 HIV associated genes from GWAS Catalog (www.ebi.ac.uk/gwas/), literature and gene-diseases database such DisGeNET (disgenet.org). We leveraged the dbSNP151 database (https://www.ncbi.nlm.nih.gov/snp/ (122)) to extract SNVs associated with these genes in the Botswana data set (**Table S5**).

### Pathways enrichment analysis and gene-gene interactions

The GeneMANIA (39) tool was used to analyse how the genes harbouring the identified variants interact in a biological network. This allowed us to obtain an enrichment of related genes within the obtained sub-network with potential biological pathways, processes, and molecular functions. Enrichment analysis was performed using Enrichr package (40,123) in R (124).

### Population diversity

#### Principal components analysis (PCA) and admixture analysis

Variants were pruned to remove those with minor allele frequency < 5%, > 2% missingness, those that deviated from Hardy-Weinberg Equilibrium (HWE p > 1.0 x 10^−5^), and those in in linkage disequilibrium (LD) r^2^ < 0.85 within 1000kb window size, incrementing with 50 bases step (--indep-pairwise 1000 50 0.15). (--indep-pairwise 1000 50 0.15). This resulted in 258,773 variants retained for assessing population diversity. For admixture analysis, we analysed the merged dataset of a total of 5,322 samples including Botswana HIV-1 positive/negative individuals and the 20 world-wide ethnic groups (**Table S7**). The ADMIXTURE (125) algorithm was used to estimate the ancestry proportions of the Botswana HIV-1 positive/negative groups. To evaluate the extent of substructure in the Botswana HIV-1 positive/negative population and whether stratification can be accounted for in the genetic association tests, PCA implemented in the **smartpca** programme in the EIGENSOFT package (17,18) was applied to the merged data set. We assessed structure between the population of Botswana and the 20 world-wide ethnic groups. Population structure and admixture were visualized by PCA plots generated using Genesis software (126) and R (124) with the pca3d package (127).

#### Distribution of genetic ancestry proportions by HIV-1 positive/negative status

The mean proportions of the three ancestries were compared between HIV-1 positive and HIV-1 negative individuals. The accurate admixture cluster was identified from model inference with lowest cross-validation (CV) error and the genome-wide admixture proportion estimations of that model inference were used as accurate genetics ancestry contribution. From these, and also basing on the population history of Southern Africa, we chose the best 3 proxy ancestral populations that had the highest genome-wide ancestry proportions from admixture analysis: Niger-Congo, Khoe-San and European.

#### Genetic distance estimated by F_ST_

Pairwise genetic distance was estimated between the Botswana population and the 20 world-wide ethnic populations using the Weir and Cockerham’s F_ST_ (128) in PLINK. A heatmap of the genetic distances was generated using package (129) in R (124).

#### Genetic relatedness and runs of homozygosity

We assessed cryptic relatedness in the population of Botswana using PLINK. Pairwise allele sharing (identity-by-descent, IBD) was determined using pi_hat threshold of 0.2 (--genome -- min 0.2). We further used PLINK to calculate homozygosity by keeping some of the default parameters while adjusting the window length and number of heterozygous SNVs allowed in the window (--homozyg-kb 150 and --homozyg-window-het 3). We compared the median lengths and segments of the runs of homozygosity (ROH) between the Botswana individuals and other world ethnic groups using Mann-Whitney U test in R (124).

## ACKNOWLEDGEMENTS

This work was supported through the sub-Saharan African Network for TB/HIV Research Excellence (SANTHE), a DELTAS Africa Initiative [grant # DEL-15-006]. The DELTAS Africa Initiative is an independent funding scheme of the African Academy of Sciences (AAS)’s Alliance for Accelerating Excellence in Science in Africa (AESA) and supported by the New Partnership for Africa’s Development Planning and Coordinating Agency (NEPAD Agency) with funding from the Wellcome Trust [grant # 107752/Z/15/Z] and the UK government. The views expressed in this publication are those of the authors and not necessarily those of AAS, NEPAD Agency, Wellcome Trust, or the UK government. The authors would also like to thank the National Research Foundation of South Africa for funding (NRF) [grant # RA171111285157/119056]. We thank the participants, investigators and key personnel of the “Host Genetics of HIV-1C Infection, Progression, and Treatment in Africa/GWAS on Determinants of HIV-1C Infection” study at Botswana Harvard AIDS Institute Partnership. All computational analysis was performed through the Centre of High Performance Computing cluster (Cape Town, South Africa).

## FUNDING

PT is a PhD student funded by the Sub-Saharan African Network for TB/HIV Research Excellence (SANTHE), a DELTAS Africa Initiative [grant # DEL-15-006].

### DECLARATION OF INTEREST

The authors declare that they have no competing interests.

## AUTHORS CONTRIBUTIONS

PKT, ERC and SG conceived and structured the content of the manuscript. PT and ERC conducted data analysis and result interpretation. PKT drafted the manuscript. PKT, ERC, WTC, DDM, CD, MML, VN, SJO, ME and SG edited the manuscript.

## Notes

### Competing Interest Statement

The authors have declared no competing interest.

