## Supplementary information for "Whole Genome Sequencing-based Characterization of Human Genome Variation and Mutation Burden in Botswana"

#### Variant Calling parameters applied to the sequence data

All the sequences passed quality control. Over 90% of the samples had a high sequence quality (at least 30 Phred score). This means that for these sequences a 99.9% accuracy in base calling was achieved. The acceptable Phred score for sequencing downstream analyses (population structure and genetic association) is at least 20. Although the sequence quality of the study samples was good, for variant calling bases with a Phred score of at least 30 were considered to ensure accurate identification of variants.

For **GATK** variant calling and filtration, we used GATK version 4.1.4.1. We performed variant calling using GATK's HaplotypeCaller to identify potential variants in each sample, then performed population joint genotyping to ensure high accuracy of calling. The variant calling followed GATK's best practices the following parameters:

```
gatk --java-options " -Xmx8g " HaplotypeCaller \
-R hg38.fasta \
-I SAMPLE.bam \
--dbsnp dbsnp151.vcf.gz \
--emit-ref-confidence GVCF \
-stand-call-conf 30 \
-O SAMPLE.g.vcf
```

The genotype VCF files were combined into a cohort file then population joint-calling was performed using the following parameters:

```
gatk --java-options " -Xmx100g" GenotypeGVCFs \
-R hg38.fasta \
--dbsnp dbsnp151.vcf.gz \
-V combine.vcf.gz \
```

```
-stand-call-conf 30.0 \
-A Coverage -A FisherStrand -A StrandOddsRatio -A MappingQualityRankSumTest \
-A QualByDepth -A RMSMappingQuality -A ReadPosRankSumTest \
--allow-old-rms-mapping-quality-annotation-data \
-O gatk.cohort.vcf.gz
```

For **BCFTOOLS** variant calling we used multiallelic calling model with the following parameters:

```
bcftools mpileup -Q30 -Ou -f hg38.fasta \
SAMPLE1.bam \
SAMPLE2.bam \
SAMPLE390.bam | bcftools call -mv -Oz -o bcf.cohort.vcf.gz
```

Raw variants called from GATK were filtered on minimum depth (DP) of 10, minimum genotype quality (GQ) of 20 and genotype call rate of 90% using VCFTOOLS. Following GATK guidelines, we filtered variants that had excess heterozygosity (> 54.69) and further filtered the data using machine learning framework implemented in GATK's Variant Quality Score Recalibration (VQSR) with the above-mentioned annotation features and the following training data sets:

**INDELs**

Mills_and_1000G_gold_standard.indels.hg38.vcf (prior=12.0)
dbsnp151.vcf (prior=2.0)

**SNVs**

hapmap_3.3.hg38.vcf (prior=15.0)
1000G_omni2.5.hg38.vcf (prior=12.0)
1000G_phase1.snps.high_confidence.hg38.vcf (prior=10.0)
dbsnp151.vcf (prior=7.0)

The training models were applied to the data using GATK's ApplyVQSR with truth sensitivity level of 99.9%.

We used the following filters on the BCFtools call set: and the following filters: depth (DP) > 10; mapping quality (MQ) > 30; variant quality (QUAL) > 20; <10% missing genotypes; <10% heterozygosity; and filtering SNP within 3bp around a gap (--SnpGap 3).

#### *Variant characterisation*

A total of 27,662,062 variants were discovered in the whole genomes of 390 individuals from Botswana (**Table S1**).

**Table S1. Total variants discovered in the genomic data of Batswana**

| **Functional class** | **Number of variants** |
|---|---|
| UTR3 | 237690 |
| UTR5 | 23702 |
| UTR5 and UTR3 | 100 |
| downstream | 169730 |
| exonic | 171348 |
| exonic and splicing | 69 |
| intergenic | 15116866 |
| intronic | 10085022 |
| ncRNA exonic | 89096 |
| ncRNA exonic and splicing | 20 |
| ncRNA intronic | 1646163 |
| ncRNA splicing | 578 |
| splicing | 2868 |
| upstream | 114776 |
| upstream and downstream | 4034 |
| **Exonic variants** | |
| nsSNV | 91994 |
| sSNV | 72040 |
| FS indel | 2964 |
| stopgain | 1635 |
| nonFS indel | 1635 |
| stoploss | 123 |
| unknown | 957 |

A total of 2,789,599 novel variants were discovered in the genomes of Batswana. The categories of these variants are presented in **Table S2**.

**Table S2.** Novel variants discovered in the genomic data of Batswana

| **Functional class** | **Number of variants** |
|---|---|
| UTR | 29904 |

| | |
|---|---|
| flanking | 30731 |
| exonic | 21665 |
| intergenic | 1461193 |
| intronic | 1066166 |
| ncRNA | 178178 |
| splicing | 1762 |
| **Exonic variants** | |
| nsSNV | 13769 |
| sSNV | 5670 |
| FS indel | 1313 |
| stopgain | 412 |
| nonFS indel | 380 |
| stoploss | 25 |
| unknown | 96 |

The distribution of novel variants was compared among four classes of minor allele counts (MAC) in **Table S3**.

**Table S3. Novel variants per MAF category**

| Functional class | MAF | | | |
|---|---|---|---|---|
| | >0-0.01 | >0.01-0.05 | >0.05 | Total |
| UTR | 28370 | 1503 | 31 | 29904 |
| flanking | 28962 | 1725 | 44 | 30731 |
| exonic | 21160 | 500 | 5 | 21665 |
| intergenic | 1374111 | 85337 | 1745 | 1461193 |
| intronic | 1007198 | 57925 | 1043 | 1066166 |
| ncRNA | 167663 | 10331 | 184 | 178178 |
| splicing | 1758 | 3 | 1 | 1762 |
| Total | 2629222 | 157324 | 3053 | 2789599 |

**Table S4. The most deleterious (damaging) variants as predicted by at least 10 databases in ANNOVAR**

| CHR | POS | SNV | REF/ALT | cDNA change | AA change | Gene | FATHHM | Siphy_29 | CADD_phred | LOF_rare* | Freq_BW | gnomAD_AFR | Freq_EU |
|---|---|---|---|---|---|---|---|---|---|---|---|---|---|
| 1 | 1184029 | rs111377341 | C/G,T | exon8:c.C979G | p.R327G | *TTLL10* | T | 11.689 | 20.6 | . | 0.033,0.005141 | 0.000309112,0.00076089 | 0,1.54851e-05 |
| 1 | 3022425 | rs3795263 | G/A,C | exon1:c.G739A | p.G247R | *ACTRT2* | D | 11.17 | 16.11 | . | 0.001289,0.003866 | 0.0441905 | 0.236254 |
| 1 | 8863242 | . | G/A,T | exon7:c.C890A | p.T297N | *ENO1* | T | 18.297 | 27.7 | stopgain | 0.001282,0.001282 | 0 | 1.55E-05 |
| 1 | 47140895 | rs61507155 | A/T,C | exon2:c.A311T | p.Y104F | *CYP4A22* | T | 5.486 | 17.08 | . | 0.035,0.001282 | 0.0645445 | 0.000325198 |
| 1 | 205844819 | rs374784539 | C/A,G | exon4:c.G568C | p.D190H | *PM20D1* | T | 19.649 | 18.79 | . | 0.001282,0.002564 | 4.76E-05 | 0 |
| 2 | 176100737 | rs200302685 | G/C,A | exon2:c.G790C | p.E264Q | *HOXD12* | D | 19.976 | 34 | . | 0.032,0.002564 | 0.000641818,0.000380337,2.3771e-05 | 0.000139336,9.28908e-05,0.000216745 |
| 2 | 195906730 | rs168192 | G/T,C | exon27:c.C4264G | p.P1422A | *DNAH7* | T | 17.963 | 17.81 | stopgain | 0.094,0.003846 | 0.049666 | 0.000372151 |
| 4 | 95841276 | rs17024795 | C/G,A | exon1:c.C1126G | p.R376G | *PDHA2* | T | 10.855 | 14.07 | . | 0.027,0.001282 | 0.0407337,0 | 0.0000309809,1.54904e-05 |
| 4 | 150599048 | rs111883007 | G/C,A | exon38:c.C6005G | p.S2002W | *LRBA* | T | 19.872 | 22.2 | FS insertion | 0.005128,0.001282 | 0 | 0 |
| 4 | 154234762 | rs61746111 | G/A,T | exon25:c.C8525A | p.P2842Q | *DCHS2* | T | 20.346 | 17.9 | FS insertion | 0.032,0.003846 | 0.0370177,0.00335395 | 0.000154895,1.54895e-05 |
| 5 | 83512337 | rs183969050 | T/C,G | c.T983G | p.F328S | *VCAN* | T | 10.479 | 18 | FS insertion | 0.013,0.001282 | 0.000356888 | 0 |
| 5 | 90652411 | rs147062294 | C/G,T | exon19:c.C3482G | p.S1161C | *ADGRV1* | T | 20.672 | 34 | FS insertion | 0.014,0.001282 | 0.0149374 | 0.000108416 |
| 6 | 43675226 | rs35480020 | G/C,A | exon5:c.C422G | p.P141R | *MRPS18A* | . | 12.36 | 17.34 | . | 0.022,0.001282 | 0,0.0353417 | 0.0000154866,0.000278759 |
| 7 | 20727068 | rs111647033 | G/A,C | exon13:c.G1319C | p.R440P | *ABCB5* | D | 15.611 | 20.1 | . | 0.026,0.002564 | 0.000523585,0.00218954 | 0,0 |
| 7 | 134538257 | rs142388608 | C/G,T | exon8:c.C805G | p.R269G | *AKR1B10* | T | 11.869 | 15.57 | FS deletion | 0.007692,0.001282 | 0.000214133,0.000118963 | 0.000139371,0.000185828 |
| 8 | 24487298 | rs146451180 | G/A,T | exon11:c.G1072A | p.V358M | *ADAM7* | T | 9.477 | 14.63 | splicing | 0.001282,0.002564 | 0.000499762 | 0.00142455 |
| 11 | 4946887 | rs2412467 | C/T,G | exon1:c.G214A | p.D72N | *OR51A4* | T | 14.304 | 18.96 | . | 0.002564,0.001282 | 0.0168363 | 0.038437 |
| 11 | 6891758 | . | A/C,T | exon1:c.T743A | p.V248D | *OR2D2* | T | 13.213 | 19.01 | . | 0.002564,0.003846 | . | . |
| 12 | 52775400 | rs149569624 | T/A,C | exon2:c.A803T | p.D268V | *KRT76* | T | 14.612 | 22.8 | . | 0.01,0.012 | 0.00497004,0.00998811 | 0,4.64626e-05 |
| 14 | 20641145 | rs148281603 | C/G,A | exon1:c.G547C | p.D183H | *OR6S1* | T | 13.924 | 19.52 | . | 0.013,0.007692 | 0.00107112,7.14082e-05 | 0,0 |
| 15 | 49972713 | rs77004004 | G/C,T | exon13:c.C1112A | p.P371H | *ATP8B4* | D | 8.864 | 14.25 | . | 0.04,0.019 | 0.0000238243,0.0132463 | 0,0 |
| 16 | 48139198 | rs113496237 | C/T,G | exon5:c.G796C | p.G266R | *ABCC12* | D | 15.741 | 19.24 | . | 0.021,0.013 | 0.0000714184,2.38061e-05 | 0,0 |
| 17 | 3292209 | rs703903 | C/T,A | exon1:c.G374T | p.R125L | *OR3A1* | T | 17.71 | 29.8 | . | 0.16,0.001282 | 0.0132319,0.295096 | 0.000077553,0.542611 |
| 20 | 35801566 | . | T/A,G | exon2:c.T44A | p.V15E | *PHF20* | T | 15.004 | 23.8 | . | 0.001282,0.001282 | . | . |

*The function of an additional rare, novel variant. CHR: chromorome, SNV: single nucleotide variant, REF/ALT: reference allele/alternative allele, AA change: amino acid change, Freq_BW: frequency in the Botswana data, gnomAD_AFR: frequency in the gnomAD African/African-American population, Freq_EU: frequency in theEuropean population.

**Table S5. Human genes previously associated with HIV-1**

| **HIV-1 associated genes** |
|---|
| A4GALT, ABCB1, ABCF2, ABO, ABTB2, AC005062.1, AC006305.1, AC009132.1, AC009229.1, AC009271.7, AC010476.2, AC010519.1, AC010998.3, AC011306.1, AC013727.1, AC018880.2, AC021573.1, AC022483.1, AC023798.16, AC023950.6, AC025614.2, AC026341.1, AC026371.1, AC034229.1, AC037459.4, AC044836.1, AC062039.1, AC063952.2, AC073475.1, AC087190.5-2, AC087854.1, AC091905.4, AC092745.2, AC092745.5, AC093801.2, AC095058.3, AC096719.1, AC097652.1, AC099795.1, AC100812.1, AC103881.1, AC104232.2, AC105916.1, AC114322.1, AC115283.1, AC116362.1, AC126407.1, ACTR3BP6, ADAM10, ADAM18, ADAMTS1, ADH5P4, AE01, AF233439.1, AGAP2, AGBL5, AJ239318.1, AJ239321.1, AKR7A2, AKT1, AL008729.1, AL031315.1, AL035246.1, AL035461.3, AL096854.1, AL136363.2, AL138752.2, AL138889.1, AL161781.1, AL353743.2, AL354694.1, AL358787.1, AL359382.2, AL390763.1, AL391500.13, AL391832.4, AL451007.2, AL451127.2, AL512329.2, AL513164.1, AL591509.5, AL671762.1, AL671883.2, AL671883.3, ALK, ALKBH8, ANKRD22, ANKRD30A, ANKRD43, ANKRD6, ANKRD9, ANXA1, AOAH, AP001021.1, AP001021.3, AP003398.2, AP003464.1, AP2M1, APOBEC3, APOBEC3B, APOBEC3G, ARF1, ARGLU1, ARHGAP32, ARHGEF12, ARHGEF19, ARPC1A, ASXL2, ATG12, ATG16L2, ATG7, ATP6V0A1, BAHD1, BCL9, BICRA, BIRC4BP, BOD1P, BRWD2, BSDC1, BTNL2, BUD13, BX323046.2, C10orf11, C10orf71, C1ORF103, C21orf96, C3ORF56, C4orf17, C6ORF1, C6orf106, C6orf15, C6orf48, C7orf58, CACNG1, CADM1, CADPS, CAPN6, CARD16, CAV2, CBS, CCBE1, CCDC134, CCL11, CCL17, CCL18, CCL2, CCL3, CCL3L1, CCL4, CCL5, CCNG1, CCNT1, CCR2, CCR5, CCRL2, CCT8L2, CD209, CD247, CD33, CD4, CDCA7L, CDSN, CHORDC1, CHRNA3, CHRNA5, CIG-5, CLDND1, CLEC18B, CLN3, CLNS1A, CMPK2, CMTM8, COG2, COG3, COG4, COLEC11, COX10-AS1, COX6A1P3, CREB5, CREBBP, CRIPAK, CRTC2, CRTC3, CSPP1, CTDP1, CTLA4, CUL5, CUX2, CX3CR1, CXCL10, CXCL11, CXCL12, CXCL9, CXCR4, CXCR6, CYP1B1-AS1, CYP3A4, CYP7B1, CYPA, CYYR1, DAB1, DARC, DC-SIGN, DCK, DDEF2, DDOST, DDR1, DDX10, DDX3X, DDX40, DDX53, DDX55, DEFB1, DEPDC5, DEPDC6, DEPTOR, DHFRP2, DHX33, DIMT1L, DISC1FP1, DMXL1, DNAJB1, DNAJC18, DNAJC27, DNAJC5B, DNAL1, DOK6, DPCR1, DPM1, DPY19L3, DPYD, DRGX, DYRK1A, DYSF, ECR777, EDNRA, EFEMP1, EFHC2, EGF, EGFR, EIF2C3, EIF3H, EPHA5, EPS8, ERCC3, ERI2, ERP27, ETF1, ETHE1, EVI5L, EXOSC3, EXOSC5, FA2H, FAM174B, FAM200B, FAM229A, FAM5B, FAM76B, FANCL, FAS, FASLG, FBN3, FBXO10, FBXO18, FBXO21, FBXW11, FCGR2A, FGD6, FGF1, FHL3, FKBP1AP4, FLII, FNTA, FOXCUT, FOXN3, FRMPD1, FUT2, FUT9, GABARAP, GALNT14, GAPVD1, GBAS, GBP1, GCK, GFRAL, GLRX3, GLTSCR1, GNPDA2, GOLPH3, GOSR2, GPC5, GPR156, GRIN2A, GRM5, GRTP1, GZMH, H3F3A, HAP1, HB1, HCG22, HCP5, HCP5HCP5, HCP5P2, HCRTR2, HEATR1, HGS, HIBCH, HIP1R, HIST1H3B, HIST1H3C, HIST1H4A, HIST1H4B, HIV-1, HLA-A, HLA-B, HLA-B57, HLA-C, HLA-DPA1, HLA-DQA1, HLA-DQB1, HLA-DRA, HLA-E, HLA-G, HMGXB3, HNRNPF, HS3ST3A1, HS6ST2, HSP90AB3P, HTATSF1, HUWE1, IDH1, IER3, IFI44, IFI6, IFIT3, IFNAR1, IFNG, IFNGR1, IGHMBP2, IGHV1-12, IGHV1OR21-1, IGHV3-13, IGHV3-52, IGHV3-53, IGSF21, IKBKG, IL10, IL10RA, IL12A, IL12B, IL13, IL2, IL2RA, IL32, IL4, IL4R, IL6, INTS7, IPO8, IQCA1L, IQUB, IRF4, ISG43, ITPKA, JAK1, JHDM1D, JUP, KAT2A, KB-67B5.12, KBTBD7, KCNIP1, KCNIP3, KCNK9, |

| |
|---|
| KCNMB3P1, KCNQ5, KDM3B, KDM4C, KDM4D, KEL, KERA, KIAA1012, KIF3C, KIF4B, KIR, KIR3DL1, KLHDC2, KLHL1, KTN1, LAPTM5, LARS, LARS2, LCP2, LEFTY1, LENG1, LINC00836, LINC00937, LINC00992, LINC01150, LINC01556, LINC01985, LINC02177, LINC02240, LINC02364, LINC02374, LINC02477, LINC02646, LINC02647, LINC02667, LINC02702, LINC02748, LNX2, LOC100129699, LOC375190, LPL, LRMDA, LRP4, LRRC58, LRRC8D, LSM3, LY6D, LYPD4, MAD2L1, MAGI1, MAP4, MBL2, MBNL2, MCM8, MDN1, MED14, MED28, MED4, MED6, MED7, MEPE, MESTP3, MGAT1, MICA, MICB, MID1IP1, MIR1275, MIR6891, MIR8074, MKI67, MKRN2, MKRN3, MMADHC, MND1, MOB1B, MOS, MPHOSPH6, MR1, NAV2, NBEA, NBPF13P, NCBP2, NCBP2-AS1, NCBP2-AS2, NCOR2, NDUFB7, NEDD9, NF2, NGLY1, NIPSNAP3B, NKG7, NLRP1, NMT1, NOS3, NOTCH4, NR0B2, NTM, NTM-AS1, NUFIP1P1, NUP107, NUP133, NUP153, NUP155, NUP160, NUP85, OAS1, OAS2, OASL, ODZ4, OTUD3, PABPC1P2, PANK1, PARD3B, PAX5, PBX1, PC, PCDH11X, PCNT, PDCD5, PDE7A, PDIA6, PHACTR1, PHF12, PHF3, PIGH, PIGK, PIGY, PIGZ, PIP5K1C, PKD1L2, PLD5, PLEKHA3, PLOD3, PM20D1, PNRC1, POLR3A, POLR3F, POU1F1, POU2F1, POU5F1, PP2672, PPIA, PPIAP33, PPIB, PPP1CB, PPP2R2A, PPP3CC, PQLC2, PRDM10, PRDM14, PRDM7, PRF1, PRKG2, PRKX, PROX1, PROX1-AS1, PSME2, PSORS1C1, PSORS1C3, PURA, RAB1B, RAB28, RAB2A, RAB6A, RAB6B, RAB6C, RABEPK, RANBP1, RANBP2, RAP1B, RAP1GAP2, RAPGEF1, RAPGEF2, RELA, RGCC, RGP1, RHOH, RICH2, RIMS4, RN7SKP199, RN7SL492P, RNA5SP407, RNA5SP408, RNF130, RNF170, RNF212, RNF26, RNF39, RNU105C, RNU6-1133P, RNU6-737P, RNU6-931P, RP11-100A16.1, RP11-707M13.1, RPL13AP15, RPL15P15, RPL19P16, RPL21P119, RPL21P126, RPL21P75, RPL23AP96, RPL28P3, RPL32P3, RPL4P5, RPS6KA2, RPSAP40, RPTN, RRAGB, RSAD2, RSL1D1, RTN2, RUNX1, RUSC2, RXRG, SAMD5, SCFD1, SCGB1D4, SCGB2A2, SDC1, SDC2, SDF1, SEC14L1, SERPINA1, SESTD1, SFT2D1, SGCD, SIGLEC17P, SIGLEC22P, SILC1, SIP1, SLC2A1, SLC2A1-AS1, SLC35F4, SLC46A1, SLC9A9, SLCO5A1, SMC4P1, SMIM12, SNHG32, SNN, SNRPEP6, SORBS3, SOX11, SOX5, SP110, SPAST, SPATS2L, SPCS3, SPDYA, SPOCK1, SPRY4-AS1, SPTAN1, SPTBN1, SSB, ST14, ST3GAL5, ST8SIA3, STAC2, STARD3NL, STAT1, STT3A, STX11, STX5, SUGCT, SUV420H1, TAOK1, TAP2, TBC1D7, TCEB3, TFAP4, TFDP2, TFE3, TGFBRAP1, THAP3, THOC2, THRAP3P1, THSD7B, TIAM2, TIMM8A, TLR7, TLR8, TLR9, TM9SF2, TMC4, TMED2, TMEFF2, TMEM132C, TMEM163, TMEM181, TMEM182, TMEM230, TMTC1, TNF, TNPO3, TNS1, TNXB, TOMM70A, TOR2A, TRAPPC1, TRIB1, TRIM10, TRIM27, TRIM5, TRIM55, TRIM58, TRMT5, TRPM6, TSBP1-AS1, TSSK3, TUBAL3, TXNL3, UBQLN4, UBQLN4P1, UGT1A11P, UGT1A12P, USP18, USP26, USP6, VANGL2, VAV2, VEGFC, VPRBP, VPS53, VWC2L, WASF5P, WDR27, WDR59, WDTC1, WNK1, WNT1, XAF1, XKR4, YPEL2, YTHDC2, ZBTB2, ZBTB20, ZBTB7C, ZCCHC7, ZDHHC19, ZFP90, ZNF12, ZNF182, ZNF354A, ZNF385D, ZNF436, ZNF512B, ZNF536, ZNF704, ZNF720, ZNF785, ZNF791, ZNF804A, ZNF831, ZNF90, ZNRD1, ZNRD1AS, ZNRD1ASP |

### *Population diversity*

#### *Population description and data acquisition*

We obtained a joint-call VCF file containing 2,504 samples from 1000 Genomes Project (The 1000 Genomes Project Consortium, 2010, 2012) and 2,428 sample from the African Genome Variation Project (Gurdasani et al., 2015) which has recently characterised the admixture across 18 ethno-linguistic groups from sub-Saharan Africa. We assembled a total of 4,932 samples. Based on initial sample description (population or country labels), we used the population ethno-linguistic information (Gudykunst and Schmidt, 1987; Michalopoulos, 2012) to categorize the obtained data per ethnic group as described in (**Table S1**) and we defined 20 world-wide ethnic groups (**Table S1**). We merged these 4,932 samples with our 390 Botswana samples resulting in a final total of 5,322 samples. The merge of our dataset and 1000 Genomes Project + African Genome Variation Project was based on overlapped SNPs using **PLINK** (Purcell et al., 2007)**.**

**Table S6. Variants data from the 1000 Genomes Project (1KGP) and the African Genome Variation Project (AGVP) used for population structure and admixture analysis.**

| Population label | Ethnic group | Population description | Total Samples |
|---|---|---|---|
| **AFR** | Afro-Asiatic-Semitic | Amhara of Ethiopia | 22 |
| | African-American | Americans of African Ancestry in SW USA (ASW) | 60 |
| | African-Caribbean | African Caribbeans of Barbado (ACB) | 96 |
| | Afro-Asiatic | Al-Gharbiyah, NA, Monufia, Kafrel-Sheikh, Mansoura, Alexandria, Dakahlia, Samanoud, Al-Buhayrah, Minya, AlSharqia, El-Mahalla all from Egypt | 99 |
| | Afro-Asiatic-Cushitic | Oromo, Somali of Ethiopia | 47 |
| | Afro-Asiatic-Omotic | Wolayta of Ethiopia | 24 |
| | Khoe-San | Khoe-San | 84 |
| | Niger-Congo-Bantu | Baganda, Banyarwanda, Barundi, Rwandese, Ugandan, Banyankole of Uganda, Bakiga, Mutanzania, Basoga, other Uganda gwas unknown, Mutooro, Batooro, Nyanjiro (Tanzania) from Uganda and Luhya in Webuye, Kenya (LWK) | 2158 |
| | Niger-Congo-Bantu-South | Zulu | 98 |
| | Niger-Congo-Volta-Niger | Esan in Nigeria (ESN), Yoruba in Ibadan, Nigeria (YRI) | 205 |
| | Niger-Congo-West | Gambian in Western Divisions in the Gambia (GWD), Mende in Sierra Leone (MSL) | 198 |
| **AMR** | Latin-American | Puerto Ricans from Puerto Rico (PUR), Colombians from Medellin, Colombia (CLM), Peruvians from Lima, Peru (PEL), Mexican Ancestry from Los Angeles USA (MXL) | 347 |
| **EUR** | European Center | British of England and Scotland (GBR) | 91 |
| | European North | Finnish of Finland (FIN) | 99 |
| | European South | Iberian Population in Spain (IBS), Toscani of Italia (TSI) | 214 |
| | European USA | Utah Residents with Northern and Western European Ancestry (CEU) | 99 |
| **EAS** | East Asian | Southern Han Chinese (CHS), Chinese Dai in Xishuangbanna, China (CDX), Kinh in Ho Chi Minh City, Vietnam (KHV), Han Chinese in Beijing, China (CHB), Japanese in Tokyo, Japan (JPT) | 504 |

| | | | |
|---|---|---|---|
| | South Asian | Punjabi from Lahore, Pakistan (PJL), Bengali from Bangladesh (BEB) | 180 |
| **SAS** | UK Indian | Sri Lankan Tamil from the UK (STU), Indian Telugu from the UK (ITU) | 204 |
| | USA Indian | Gujarati Indian from Houston, Texas (GIH) | 103 |
| **Total** | | | 4,932 |
